# A brain-inspired object-based attention network for multi-object recognition and visual reasoning

**DOI:** 10.1101/2022.04.02.486850

**Authors:** Hossein Adeli, Seoyoung Ahn, Gregory J. Zelinsky

## Abstract

The visual system uses sequences of selective glimpses to objects to support goal-directed behavior, but how is this attention control learned? Here we present an encoder-decoder model inspired by the interacting bottom-up and top-down visual pathways making up the recognition-attention system in the brain. At every iteration, a new glimpse is taken from the image and is processed through the “what” encoder, a hierarchy of feedforward, recurrent, and capsule layers, to obtain an object-centric (object-file) representation. This representation feeds to the “where” decoder, where the evolving recurrent representation provides top-down attentional modulation to plan subsequent glimpses and impact routing in the encoder. We demonstrate how the attention mechanism significantly improves the accuracy of classifying highly overlapping digits. In a visual reasoning task requiring comparison of two objects, our model achieves near-perfect accuracy and significantly outperforms larger models in generalizing to unseen stimuli. Our work demonstrates the benefits of object-based attention mechanisms taking sequential glimpses of objects.

## 1 Introduction

Objects are the units through which we interact with the world and perform tasks [70, 60, 67]. Consider the simple task of searching for an object in a cluttered environment (e.g. a paper clip in a busy drawer). This task requires one to sequentially move spatial attention to select objects and to use object-based attention to route and bind the features of these objects so that they can be recognized as a target or non-target. The role of the attention mechanism in this process is to make the objects in the sequence the figure, in order to facilitate interacting with the object and performing the task [68, 86]. Works have also suggested that object-based attention is important for achieving human-level performance in accuracy and generalization [26, 67]. However, in contrast to spatial [14, 1] or feature-based attention [52, 50], where attention studies can focus on simple, well-defined visual features (e.g., location, color, etc.), the units of selection for object-based attention [5, 18, 49] are entities consisting of complex spatial properties (shapes) and distributed feature representations. Moreover, perception is akin to hypothesis testing, where a top-down modulation reflecting object hypotheses and goals affect object selection and bottom-up processing. Leveraging recent developments in deep learning [47], here we model a general-purpose object-based attention mechanism, requiring solutions to three sub-tasks: (a) solving the binding problem, grouping features of each object, using object-centric representation learning (b) capturing the interaction between a (largely) bottom-up mechanism for recognition and a top-down mechanism for attention planning using encoder-decoder models, and (c) learning to sequentially sample objects using end-to-end training.

To solve the object binding problem [78], recent deep neural networks (DNNs) have been proposed that create object-centric representations of the entities in a scene by spatially segregating the features from the background and dynamically grouping those features in spatial and representational domains [29]. This segregation and grouping relies on (bottom-up) part-whole matching and Gestalt processes interacting with (top-down) objectness priors and knowledge of object categories [29, 86, 88]. The representation of different objects are stored in separate “slots” [29, 28, 8, 51, 21], realizing the cognitive concept of “object files” [40, 27]. A recent development in this domain, Capsule Networks (CapsNets) [66, 35], attempt to represent scenes as parse trees. Capsules in different layers represent visual entities at different levels of object granularity in the image, from small object parts in the lower levels to whole objects at the top level. Capsules provide an encapsulation and grouping of object representations that has shown benefits for performing object-based and class-conditioned reasoning for downstream tasks (e.g. deflecting adversarial attacks; [64]). Object-centric representations have great potential for modeling visual perception (e.g. for predicting global shape processing behavior in humans [17]), but to our knowledge, have not yet been studied in the context of planning attention glimpses.

An object-centric perspective requires an integrated attention-recognition mechanism that is nicely embodied by the interacting structures organized along the ventral and dorsal visual pathways in the brain [79, 80]. The ventral “what” pathway is involved in feature processing and recognizing objects and scenes [22, 79]. Core object recognition refers to the rapid recognition of briefly presented objects (100ms) [84] and is believed to be carried out primarily in the initial feedforward processing pass in the ventral pathway [15]. Modeling works support this claim by showing that neural activation along this pathway during core object recognition can be predicted [9, 31, 10] using feedforward Convolutional Neural Networks (CNNs [48, 47]) trained on the same recognition tasks. However, difficult recognition tasks require recurrent and feedback connections beyond the feedforward pass [42, 41, 71, 89]. The dorsal “where” pathway is believed to be involved in the spatial prioritization of visual inputs and the guidance of actions to objects [6, 77]. Here we take inspiration from the role played by dorsal structures in providing attentional modulation of ventral pathway activity, as shown in previous work [12, 6]. The attention signal generated in the dorsal pathway prioritizes and modulates the routing of the visual inputs in the ventral pathway for the purpose of improving object classification success and better performing the task. The repetition of this process imposes seriality on behavior when confident classification decisions are needed, making visual perception a sequential process. Previous neuro-cognitive modeling work on an integrated attention-recognition mechanism [12] have been largely hand designed and applied only to simpler stimuli, limitations that could be better addressed with a learning-based approach.

Our model of object-based attention employs the general structure of autoencoders (and more generally encoder-decoders). This class of models learn to encode the sensory input into a compact representation that captures the important aspects of the input, and then decode that representation to reconstruct the sensory input (or translate it into another modality, e.g. image to text) [45, 90]. Notably the DRAW [30] architecture introduced a sequential spatial attention mechanism to Variational Autoencoders (VAE, [45]) and showed that the model can learn to iteratively glimpse different parts of the input images and reconstruct them. Our premise is that the encoder-decoder framework loosely maps onto interactions between the dorsal and ventral processing in the human brain’s attention-recognition system. We believe core object recognition along the ventral pathway can be mapped to encoder processing. Decoder processing maps onto dorsal pathway, from which originates the top-down attention signal that modulates ventral activity. These encoder and decoder steps are taken iteratively, creating a repeating cycle of prioritization and selection. Within this framework, we present OCRA, an **O**bject-**C**entric **R**ecurrent **A**ttention model that combines recurrent glimpse-based attention and object-centric representation learning. Like a CapsNet, it performs encapsulation of features to structure the higher level representations for object processing. However, we place this structure within the aforementioned encoder-decoder model with recurrent attention, thereby enabling integration of structured information across multiple attentional glimpses. We show that capsule-based binding of object features and grouping is effective in the sequential detection of multiple objects. It is also very effective in performing visual reasoning tasks (judging whether two randomly generated shapes are the same or different) and on a challenging generalization task where the model is tested on stimuli that are different from the training set.

## 2 Results

Object-based attention mechanisms endow a model with the ability to effectively segregate and represent different objects in order to perform a task. To test this in OCRA we used multi-object recognition and visual reasoning tasks. First, the model’s ability to segregate the figure objects from background clutter is tested on the MultiMNIST-cluttered dataset[3]. Second, we test the model’s recognition performance when objects are highly overlapped (and thus occluded) using the MultiMNIST task [66]. We also examine the model’s performance on learning visual reasoning using a similar paradigm as the Task 1 in the Synthetic Visual Reasoning Test (SVRT) [23], where the model must detect whether two randomly generated objects in a given scene are the same or different. All model accuracy results reported in this section are averages of 5 runs (see 4.5 for training details and hyperparameter selection for different experiments). For recognition tasks, accuracy is measured on the image level, meaning that the response is correct only if both objects in an image are recognized correctly.

The architecture for OCRA is shown in Fig. 1. Building on the DRAW architecture [30], the “attention window” in OCRA is a grid of filters applied to a variable-sized area of the image. However, because the number of filters covering the attention window remains constant, as the window gets bigger it samples increasingly low-resolution information, creating a tradeoff between the size of the attention window and the resolution of information extracted. This property is aligned with “zoom lens” theories of human attention [20, 55], that similarly propose a tradeoff between resolution and a variable-sized attention process that can be broadly or narrowly allocated to an input (See Fig. 5 for a visual illustration of the attention mechanism). The original DRAW model [30] was formulated as an autoregressive VAE, as it was trained for stepwise self-supervised reconstruction. In contrast, our formulation uses a deterministic encoder-decoder approach. We train OCRA to predict both the category classification, based on object-centric capsules, and an image reconstruction, based on decoder output. Refer to the method section 4 for a detailed description of the model architecture and mechanisms.

**Figure 1:**
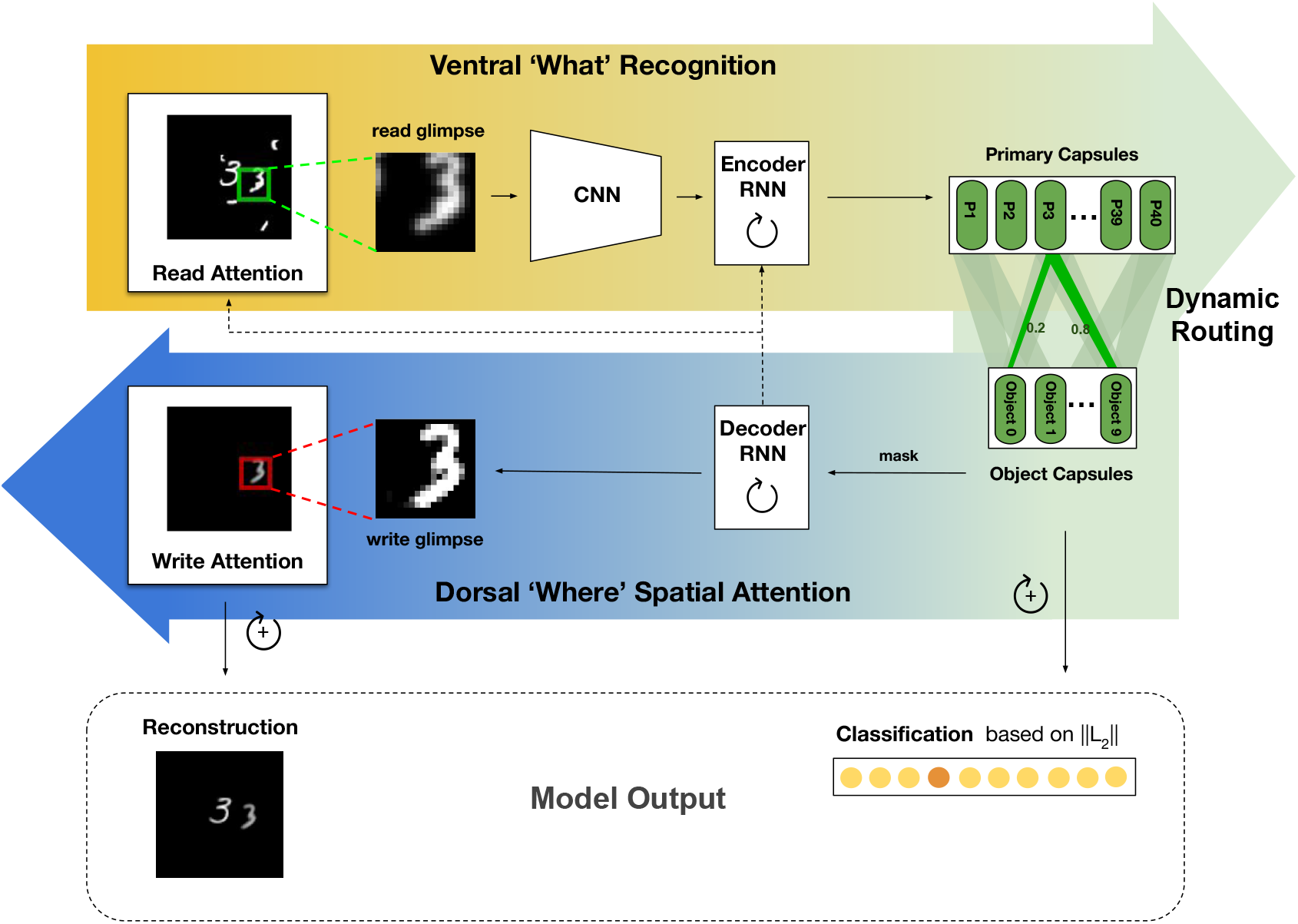
OCRA architecture. At each timestep of ventral processing, the encoder inputs a new read glimpse that is taken from the image using an attention window specified by the decoder RNN. The read glimpse is first processed using a small CNN network to obtain features that are then input to a recurrent layer. The primary capsules are read from this recurrent activation using a linear mapping, and are dynamically routed through agreement to the class capsules. The size of the the class capsules determines the evidence for a class in the selected glimpse. For the dorsal processing, the class (digit) capsules are then masked to only route the most active capsule to the decoder RNN where the representations are maintained over timesteps. This representation is used to plan attentional glimpses for both the reading operation–what gets routed in the ventral pathway, and also using a similar mechanism for reconstructing the image, by generating where and what to write to the canvas. The connection from decoder RNN to encdoer RNN allows the ongoing recurrent representation to also further modulate routing in the ventral pathway. We only use this connection for the visual reasoning task. This sequential process is iterated for multiple steps, after which the cumulative size of the class capsules determines the final classification. Both the class capsules and the cumulative canvas are used for training.

### 2.1 Objects in Clutter

Recognizing an object in a noisy environment requires the grouping and binding of the object’s features and their segregation from the noisy background and potentially other objects. We tested whether our model could learn to bind and segregate objects using the MultiMNIST-cluttered task. The stimuli for this task are generated by placing two random digits and multiple object-like distractor pieces on a canvas (see 4.1).

Fig. 2A shows OCRA’s ability to recognize the two digits over five timesteps for a sample image from this task. The top row shows the size and location of the attention window for each step. Note that OCRA makes its initial glimpse (left column) large (and therefore spatially biased to the center) so as to cover most of the image, presumably to obtain its version of scene gist. The second row shows the information being read from each attention glimpse, in the image dimension. The blurring observed over steps 1 to 3 illustrate the aforementioned zoom-lens relationship between the size of the glimpse window and its resolution, owed to OCRA’s attention having a constant number of filters. We define a reconstruction mask that is the averaged sum of all the read glimpses converted into image dimensions (average of Fig. 2A middle row). This mask would specify the image areas that the model had glimpsed by the end of each trial and is used during training to only penalize the model for not reconstructing those areas. In other words, this mask effectively focuses the loss so that the model is accountable for reconstructing only the areas where it had glimpsed, allowing the model to be selective with its glimpses and write operations and avoid glimpsing at distractors. The most active capsule that is routed to the decoder at each step, and its length, are provided at the bottom of the figure.

**Figure 2:**
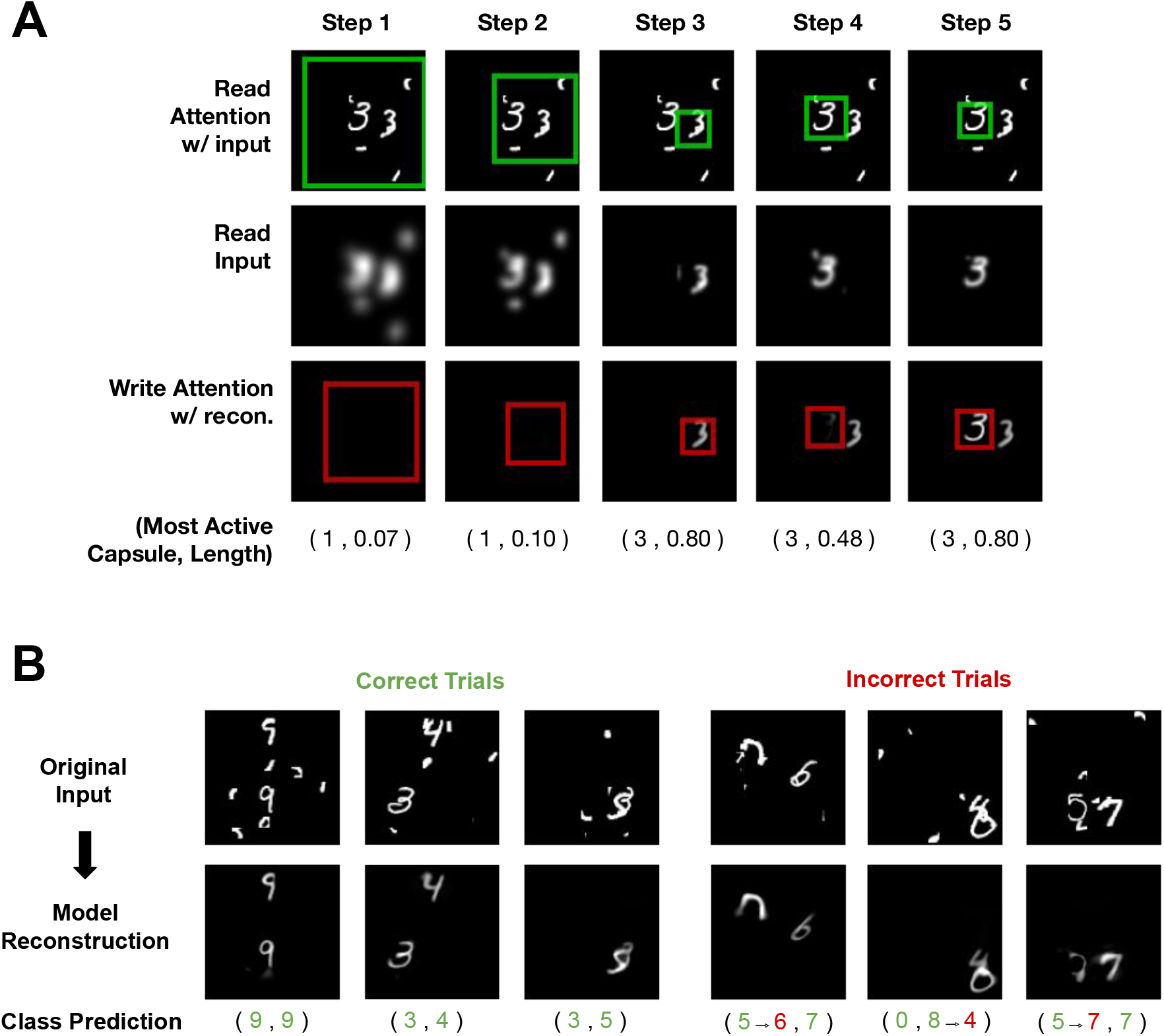
**A)** Stepwise process of moving the glimpse over a sample MultiMNIST-cluttered image. The read attention windows are overlaid as green boxes on the original image in the first row. The read glimpse are shown in the second row and the bottom row shows the cumulative canvas and the location of write attention as a red box. The most active capsule and the corresponding capsule lengths are displayed at the bottom for each step. **B)** Sample classification and reconstruction predictions for OCRA on MultiMNIST-cluttered task. The green digits show the ground truth and correct model predictions and erroneous predictions from the model are in red.

With only classification supervision, OCRA learns to take sequential glimpses of the image at different scales. Additionally, it learns to write these glimpses to the canvas only when it is confident of the digit classification. As Fig. 2A shows, this interacting attention-recognition process enables OCRA to recognize digits embedded in considerable noise. Note also that the model learns that recognition requires the sequential disengagement of attention on one object and the movement of the glimpse window to the other object so that it too can be recognized, as illustrated by the object-centric additions to the canvas in the figure. Note that this serial behavior parallels the fact that human object recognition is also serial, at least for objects complex enough to require a feature binding [78]. Fig. 2B shows more examples of model predictions, with correct responses on the left side and errors on the right. Most errors by OCRA on this task are due to the digits overlapping with each other or with noise pieces in ways that change their appearance from the underlying ground truth. Supplemental Fig. S1 shows the glimpse-based reconstruction for few sample images with correct model predictions.

OCRA performed the MultiMNIST-cluttered task with 94% accuracy, comparable to other glimpse-based models [54, 3]. While this may reflect the value of inclusion of a glimpse-based attention mechanism, first, OCRA’s attention mechanism is differentiable and therefore easy to train, in contrast to the reinforcement learning based mechanisms employed in earlier work. Another notable difference is the use of a context network in [3] that inputs the whole image to the model in a separate pathway to plan attention selection, thereby separating it from the recognition pathway. OCRA has one pathway and strings together glimpses from multiple steps to have an integrated recognition and attention planning mechanism. Our model does this through its “zoom lens” attention processes that can switch between local and global scales as needed. OCRA’s early glimpses are taken from the whole image, providing the model with a low resolution gist description (Fig. 2A, middle row, left) that allows it to plan future glimpses. Reflecting its object-centric design, OCRA’s later glimpses are focused on individual objects to mediate the recognition and reconstruction processes.

### 2.2 Overlapping Objects

Objects commonly occlude each other, and to study the attention-recognition system under these conditions we used the MultiMNIST task. This task involves recognizing digits having an average of 80 percent overlap between their two bounding boxes. Fig. 3A shows OCRA’s glimpse behavior on a sample image. The model starts with a more global glimpse but then moves its attention window, first to one object and then to the other, recognizing and reconstructing each sequentially. The most active capsule at each processing timestep is indicated by the digits below the second row. The gradual spreading of the reconstruction shown in the bottom row is consistent with the spreading of attention within an object, a pattern hypothesized by object-based models of attention [39, 18]. However, the gradual reconstruction does not show a clean cut between the two objects in the cumulative canvas, which we believe is due to the model having only seen the objects in these highly occluded images in this task (and never in isolation) and not being provided with object segmentations (only the classification ground truth). Fig. 3B shows samples of the model making correct and incorrect predictions, and the resulting reconstructions. Table 1 shows OCRA’s overall accuracy in this task (whether both objects in the image are correctly classified) compared to the CapsNet model [66] where the MultiMNIST task was originally introduced. Our model with 3 glimpses significantly outperforms the CapsNet model while having only one third the number of parameters. Finally, we observed a clear effect of the number of timesteps on model performance, with the error rate dropping from 3 timesteps to 10 timesteps. Just as human recognition benefits from longer attention sampling when discriminations are difficult, so too does OCRA’s performance.

**Table 1:**
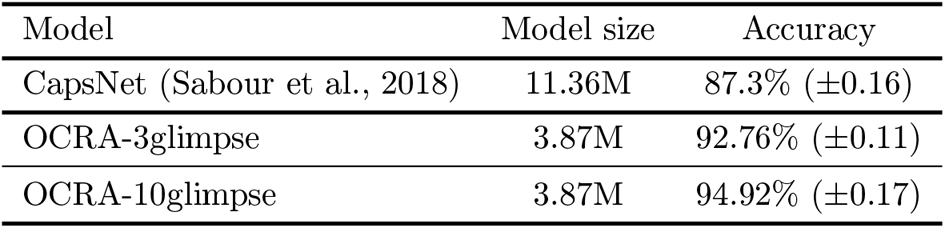
MultiMNIST classification accuracy

**Figure 3:**
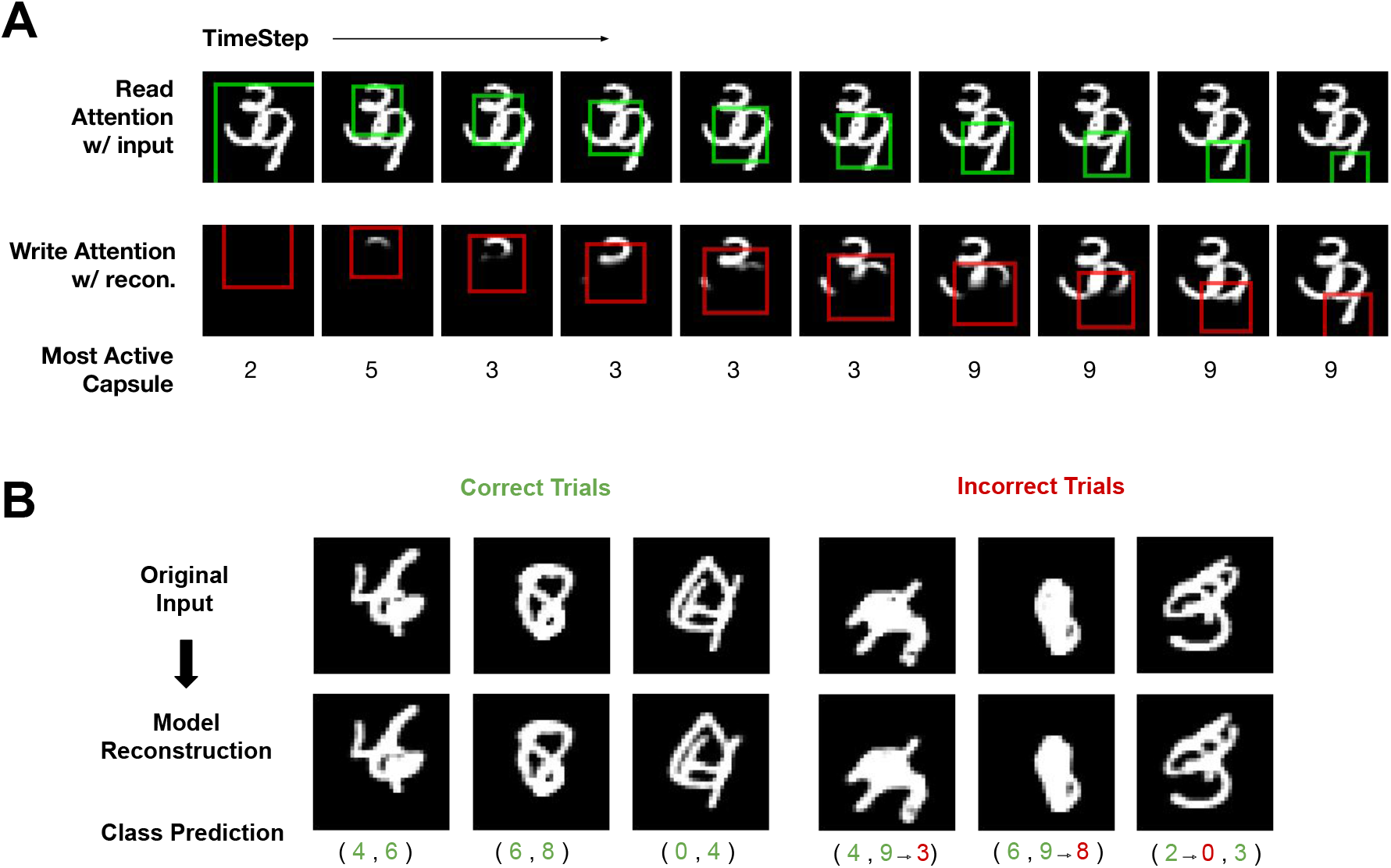
**A)** Stepwise process of moving the glimpse over a sample MultiMNIST image. The read attention windows are overlaid as green boxes on the original image in the first row. The read glimpse are shown in the second row and the bottom row shows the cumulative canvas and the location of write attention as a red box. The most active capsule and the corresponding capsule lengths are displayed at the bottom for each step. **B)** Sample classification and reconstruction predictions for OCRA on MultiMNIST task.The green digits show the ground truth and correct model predictions and erroneous predictions from the model are in red.

To more clearly understand the model dynamics, we conducted experiments starting from the OCRA-3glimpse model and ablated different model components. Namely, we conducted separate ablations for the object-centric representation, the sequential glimpse mechanism, and the model’s recurrent processing, to measure their impact on model performance. Results are shown in Table 2.

**Table 2:** Ablation study results on MultiMNIST classification

| Model | # of Glimpses | Capsules | Model size | Accuracy |
| --- | --- | --- | --- | --- |
| OCRA (proposed) | 3 | Yes | (3.87M) | 92.76% ($\pm 0.11$) |
| OCRA-Recurrent | No | Yes | (8.58M) | 91.02% ($\pm 0.14$) |
| OCRA-Feedforward | No | Yes | (6.47M) | 89.37% ($\pm 0.10$) |
| OCRA-no Capsules | 3 | No | (3.87M) | 91.96% ($\pm 0.13$) |

#### The role of recurrence and attention glimpse mechanisms

In the first ablation experiment we asked how OCRA’s performance compares to a recurrent model that uses object-centric representation but lacks an ability to obtain glimpse samples. OCRA-Recurrent (Table 2) performs multi-step processing on the input image using the recurrence in its encoder and decoder RNNs but cannot glimpse at specific locations. The model therefore receives the entire image as input at each processing step, which also requires it to have more parameters (36 *×* 36 pixel input compared to 18 *×* 18 pixel glimpse). We then trained this model using three timesteps to make it comparable to OCRA-3glimpse. As shown in Table 2, accuracy for OCRA-Recurrent is lower than for OCRA-3glimpse, highlighting the important of the glimpse mechanism in our model performance. We next removed the recurrence mechanism completely from OCRA, leaving a model that makes one feedforward pass with the full resolution image as its input. This Feedforward model then binds features for each object in separate category capsules before feeding them to the decoder (without masking the object capsules) to reconstruct the whole image at once. Table 2 shows that this Feedforward model is not only less accurate than OCRA, it is also less accurate than OCRA-Recurrent. This comparison broadly supports previous work arguing for a role of recurrent dynamics in assisting recognition tasks involving high degrees of occlusion [71, 89]. This benefit of recurrence on object recognition is hypothesized to be due to recurrence providing more computational depth [57, 82, 69] and leveraging contextualized iterative computations [82]. Taken together, while our results show that recurrent computation can improve accuracy in challenging recognition tasks with high occlusion, pairing recurrence with an attention glimpse mechanism is particularly effective in improving performance.

#### The role of object-centric representation

To examine the impact of an object-centric representation on model performance we replaced the capsule architecture with two fully-connected layers (the same size as two capsule layers; 320 and 160 units) followed by a classification readout. This new model, OCRA-no Capsules, iteratively processes the image and its classification scores are a linear readout from the second fully connected layer representation at each time step, which are then combined across timesteps to make the final classification decision. As shown in Table 2, accuracy for OCRA-no Capsules was lower than for OCRA, highlighting the effect of object-centric representation on performance in this task. However, this effect was relatively small compared to the ablations of recurrent attention, suggesting that a recurrent model can to an extent compensate for the reduced information encapsulation that results from removal of the capsule architecture. The model has a glimpse mechanism and is free to route information globally, thereby potentially reducing the benefit derived from feature encapsulation on this task.

### 2.3 Visual Reasoning

The ability to reason over visual entities and the relations between them is an important cognitive ability in humans and other animals [58, 32] with perhaps the ability to judge whether two objects are the same or different being the most fundamental form of visual reasoning. Fleuret et al. [23] proposed the Synthetic Visual Reasoning Test (SVRT) as a benchmark for testing this capability in AI systems. Here our focus is on task 1 in this benchmark (see examples in Fig. 4A), which tests a model’s ability to judge whether two randomly generated objects in an image are the same or different. While behavioral results showed that humans were able to perform this task easily, early CNN models applied to this task failed to reach high levels of accuracy, thereby exposing a shortcoming [74, 43]. However, later works found that deeper CNN models (with more layers and residual connections) such as the ResNet [33] family of models can perform this task at near perfect accuracy (>99%) while shallower models such as AlexNet [46] stay at chance level [53, 24]. This comparison identifies computational depth as an important factor in learning this task.

**Figure 4:**
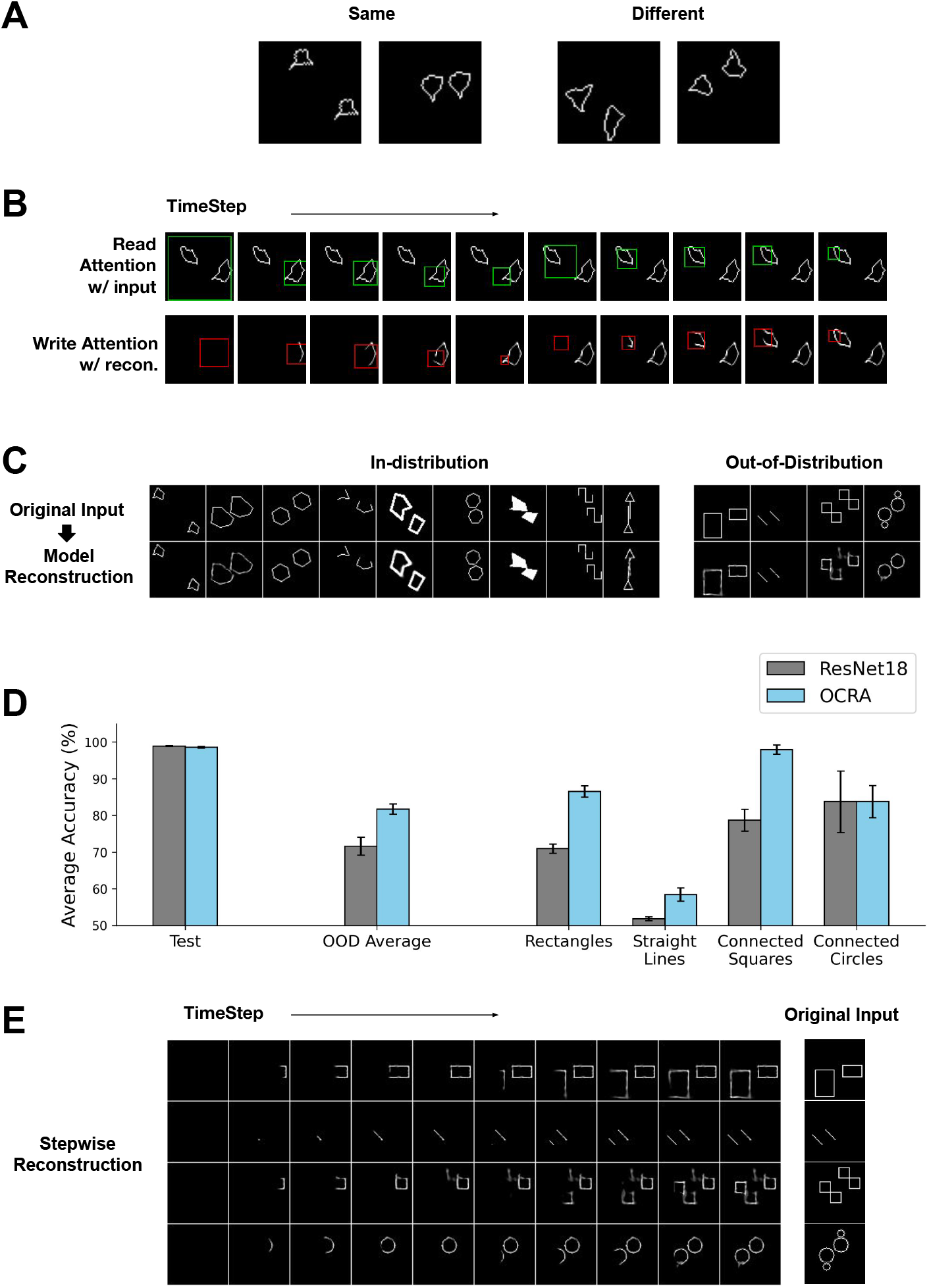
OCRA performing visual reasoning tasks. **A)** Sample images from SVRT task 1, which requires a same or different response. **B)** Stepwise visualization on a sample testing image on SVRT task 1. The top row shows the model’s read attention window in green at each time step. The bottom row shows the cumulative canvas with each write attention window in red. **C)** Sample images and model reconstruction on the OOD generalization task with each column corresponding to a different object shape. Left: in-distribution testing dataset, Right: out-of-distribution testing dataset **D)** Average model accuracy for both ResNet18 (gray) and OCRA (cyan) on the original test set and the OOD test set on average, by OOD shape type. Both models achieve near perfect accuracy on the original test set, but OCRA significantly outperforms the ResNet18 on most of the generalization tasks. Error bars show standard error **E)** The sequential processing of four OOD sample images, with each row corresponding to a different OOD shape. The original images are shown on the right and the stepwise cumulative reconstructions are shown on the left, which clearly illustrate object-based behavior

Interestingly, alternative approaches have also achieved high levels of accuracy on this task despite using smaller networks. For example, siamese networks, where two networks using shared parameters are applied to the two objects in the scene, are able to learn this task [43]. Placing the objects in different image channels of the input can improve the models’ performance on this task as well [73]. In the context of attention, these approaches are assuming a pre-attentive binding of features to objects by feeding the objects separately to each network (the case with siamese networks) or by providing them in separate channels. This assumption is broadly consistent with object-based attention, but it avoids the role of attention in solving the binding problem, which has long been considered a fundamental function of attention [78]. Consistent with OCRA and as argued by [72] (also see [65]) a more general approach would be to model the object-based attention process itself and to detect and bind the features of the objects through application of sequential processing (see [81] for an alternative approach for studying the role of bottom-up feature and spatial attention but focused on training efficiency on this task). Additionally, greater computational depth can be achieved through more recurrent sequential processing rather than a greater number of layers [69].

As shown in Fig. 4B (top row), the model starts with a global glimpse but then immediately moving to a serial attention allocation, moving its glimpse window first to one object and then to the other. Expounding on this serial behavior, the first-attended object is represented by one object capsule and this object-centric representation is maintained in the encoder-decoder RNNs across timesteps. When the model glimpses the second object, the recurrent representation would then bias the model to route information from the second object to one of two judgment capsule slots (same or different). This biasing is realized by a feedback connection from the decoder RNN to the encoder RNN that we added to OCRA for this task. The encoder RNN therefore uses feedfoward, recurrent and feedback inputs to update its activity as each timestep. The model is trained using the classification and reconstruction losses similar to the previous tasks. Over training the model learns to route and reconstruct the second glimpsed object to the first classification capsule if its features match those of the first object (which are being maintained through recurrent processing) and to the second classification capsule if the features are different. Note that we do not have explicit working memory modules in our model but the connection loops and the local recurrent connections do realize that function by maintaining the features of the first glimpsed object. In addition to capsules dedicated to the two objects, we also anticipated the need for a gist capsule (used most often for the initial glimpse) and perhaps a temporary object representation that forms when the glimpse moves between objects, bringing the total number of capsules used in the representational bottleneck to four. The bottom row of Fig. 4B shows the cumulative write canvas with the location and extent of the write attention for each of the 10 timesteps. Note again that OCRA does not just reconstruct the two objects serially, its attention also gradually spreads within an object [39, 18].

Table 3 shows model accuracy on this task. OCRA achieved the same near perfect level of accuracy (99.1%) as a ResNet18 model on this task. We then performed two ablation studies to examine the effect of the feedback connection and the object-centric representation bottleneck. The model without capsules is constructed similarly to the previous section; capsule layers were replaced by fully-connected layers and a classification readout. For the no-feedback model, we simply removed the feedback connection between the decoder RNN and the encoder RNN. We found that ablating either of these components from OCRA significantly impacted its accuracy (Table 3). This shows that OCRA performs this task well by learning to leverage both an object-centric representation and a feedback mechanism for top-down biasing in order to modulate bottom-up information routing.

**Table 3:** Ablation study results on visual reasoning

| Model | Accuracy |
| --- | --- |
| ResNet18 | 99.1% ($\pm 0.43$) |
| OCRA | 99.14% ($\pm 0.38$) |
| OCRA-no Capsules | 91.38% ($\pm 2.79$) |
| OCRA-no Feedback | 92.10% ($\pm 1.81$) |

#### Generalization to out-of-distribution stimuli

As discussed earlier, deeper CNNs can solve the same-different task in the SVRT benchmark. However, recent work revealed a major weakness [62]. These authors argued that the true test of whether a model learned the concept of “sameness” is to show that it can generalize to other displays where the pixel distribution of the generated objects is different from the pixel distribution in the training set (e.g. different shapes). Models evaluated on this dataset would therefore be asked to generalize to out of distribution (OOD) stimuli, something that humans do well. We tested OCRA and ResNet18 on a task adopted and modified from [62]. The training set includes a collection of nine different types of stimuli, as shown in Fig. 4C (left, top row). This was done to create a diverse training set to allow the models to learn a rich feature representation for performing this task. We then evaluated the models on the original test set and the OOD generalization test set, which is a collection of four types of stimuli with different shapes from the training set (examples shown in Fig. 4C, right top row).

Fig. 4C bottom row shows examples of OCRA reconstructions for some in-distribution test samples on the left (see Supplemental Fig. S2 for the cumulative canvas obtained over successive glimpses of attention for these samples) and OOD test samples on the right. As shown in Fig. 4D, OCRA and the ResNet model both achieved near perfect accuracy on the in-distribution test set for this task (leftmost bars). We found a very different result on the OOD generalization test, where OCRA significantly outperformed the ResNet model; Fig. 4D shows the average accuracy over the OOD dataset and comparisons divided by the OOD test shape. These results confirm previous work [62] showing that ResNet models critically struggle with same-different judgment generalization. However, OCRA’s sequential and object-centric processing of the image generalizes to the OOD samples, as evident in the cumulative canvas for the four examples shown in Fig. 4E, enabling significantly better performance. Lastly, note that OCRA’s reconstruction quality for the OOD images is not as good as for the original test images (comparing Fig. 4C left and right bottom rows), showing that the model still struggles with representing these stimuli and that there is room for improvement.

## 3 Discussion

Object-based attention endows humans with the ability to perform an unparalleled diversity of tasks, and we took inspiration from this mechanism in building OCRA. Its object-based sequential processing and recurrent computations, combined with its cognitively plausible (zoom-lens) glimpse mechanism, gives OCRA processing depth comparable to larger networks. OCRA’s two-pathway architecture also took inspiration from the ventral and dorsal pathways in the primate brain. Its behavior reflects an interaction between bottom-up an top-down processes such that feedforward processing is enriched and guided by current top-down hypotheses. The results show that the synthesis of these principles can lead to a model having improved robustness and generalizability, noting that we demonstrated this under challenging conditions where a mechanism for parsing the scene into entities and a recurrent process for iterative attentional sampling might prove beneficial. We also observed in the model behavior another hallmark of object-based attention, the gradual filling in of the objects [39], although for now this remains only a qualitative observation.

While many works at the intersection of machine learning and cognitive neuroscience have focused on convolutional and recurrent neural networks, encoder-decoder methods have the potential to be very useful in modeling behavior and brain responses. The generative nature of this method makes it suitable for modeling different top-down modulations and feedback processing. To date, these models have been used to study effects of top-down feedback in ventral pathway [76, 2] and to model predictive coding [37], mental imagery [7] and continual learning [83]. More generally, these models can also be used for representation learning, where they can be trained using self-supervised methods to generate the visual input. In doing this, the bottleneck representation becomes a more compact representation of the image, which through the introduction of additional constraints can be disentangled and made interpretable [75, 34]. In our work we used this framework to model, not only ventral pathway processing, but also its interaction with dorsal feedback addressing the relative paucity of work in modeling the dorsal pathway (but see [4] for a dual-stream architecture interpretation) compared to the ventral pathway. Our work is therefore timely, and will hopefully help to close what is a gap in the cognitive modeling literature relating attention and recognition.

OCRA employed a glimpse-based (“hard”) attention mechanism as this approach has the promise of making recognition and reasoning models more accurate, efficient and interpretable as they would only have to focus processing resources on smaller and relevant areas of the image [54, 3, 11, 19]. An alternative approach is for the top-down processing to create spatial masks to route object-specific information from one layer to the next [49]. Another approach toward modeling top-down attention has focused on “soft”-attention models that capture certain aspects of feature-based attention [14, 52], where top-down modulations weight the incoming representations based on task relevance [90, 56]. In this vein, transformer-based attention mechanisms [85] have recently been used to create models with an integrated process of soft attention sampling and recognition [92, 38], leading to better performance on adversarial images. In these models, the ongoing state representation produces queries that are matched to keys from the bottom-up processing to create attention weightings for new sampling of bottom-up values and the updating of the state representation. We believe that combining these approaches with the proposed glimpse based mechanism helps to build a more robust model, one that not only learns an optimal policy for making attention glimpses but also to bias feature processing along the processing hierarchy via top-down modulation in order to reflect current object hypotheses.

The discovery of objects in the visual input, and representing them in ways that are conducive to performing downstream tasks, has become an attractive area of research for understanding the relationship between deep learning (connectionist) approaches and human like symbolic and object-based reasoning [25, 29]. In these models, a first stage of generating segmented representations of the objects in a scene is followed by a second stage of modeling the interaction between these objects, either through a key-query attention mechanism (similar to transformers) [27, 16] or by graph neural networks [63]. These object-centric models provide the necessary representational capabilities to realize object-files and make broad connections to object-based effects on behavior [60, 27]. These approaches allow a model to work and learn at an object level, and should be particularly useful in modeling the varied aspects of object-based attention [61, 87, 49, 26, 13]. In return, we believe that this object-centric perspective can facilitate translation of ideas and inductive biases from the object-based attention literature, and cognitive science more generally, to build more robust and generalizable AI systems.

## 4 Materials and Methods

### 4.1 Stimuli generation

#### MultiMNIST-cluttered dataset

We generated the MultiMNIST-cluttered dataset, similar to the cluttered translated MNIST dataset from [54]. For each image, 2 digits and 6 digit-like clutter pieces are placed in random locations on a 100 *×* 100 blank canvas. All digits were sampled from the original MNIST dataset[48] and the two digits in each image could be from the same or different categories. Clutter pieces were generated from other MNIST images by randomly cropping 8 *×* 8 patches. We generated 180K images for training and 30K for testing, ensuring to maintain the same MNIST training/testing separation.

#### MultiMNIST Dataset

We generated the MultiMNIST dataset following the method from [66]. Each image in this dataset contains two overlapping digits sampled from different classes of the MNIST hand-written digits dataset[48] (size 28 *×* 28 pixels). After the two digits are overlaid, each is shifted randomly up to 4 pixels in horizontal and vertical directions, resulting in images of size 36 *×* 36 pixels with on average 80% overlap between the digit bounding boxes. We generated 3M images for training, and 500K images for testing, and ensured that the training/testing sets were generated from the corresponding MNIST sets (i.e., the training set in MultiMNIST was only generated from the training set in MNIST).

#### SVRT task 1

We generated the SVRT stimuli for task 1 directly using the code from [23]. We generated 60K images for training and validation and 10K images for testing. The images are 64×64 pixels. The stimuli for the generalization task were generated using the code modified from [62]. For each of the nine types we generated 6K images for training (54K in total) and 1.2k for testing (10.8K in total). We also generated 5.4k images for the four OOD shape types that were only used for testing.

### 4.2 OCRA Architecture

The OCRA architecture is shown in Fig. 1. An overview of the OCRA components, loss functions with pseudocodes and implementation details are provided in this section. A pytorch[59] implementation of OCRA with additional details and results are provided in these repositories: github.com/hosseinadeli/OCRA and github.com/hosseinadeli/OCRA_samediff.

#### Pseudocode 1

OCRA Architecture Overview

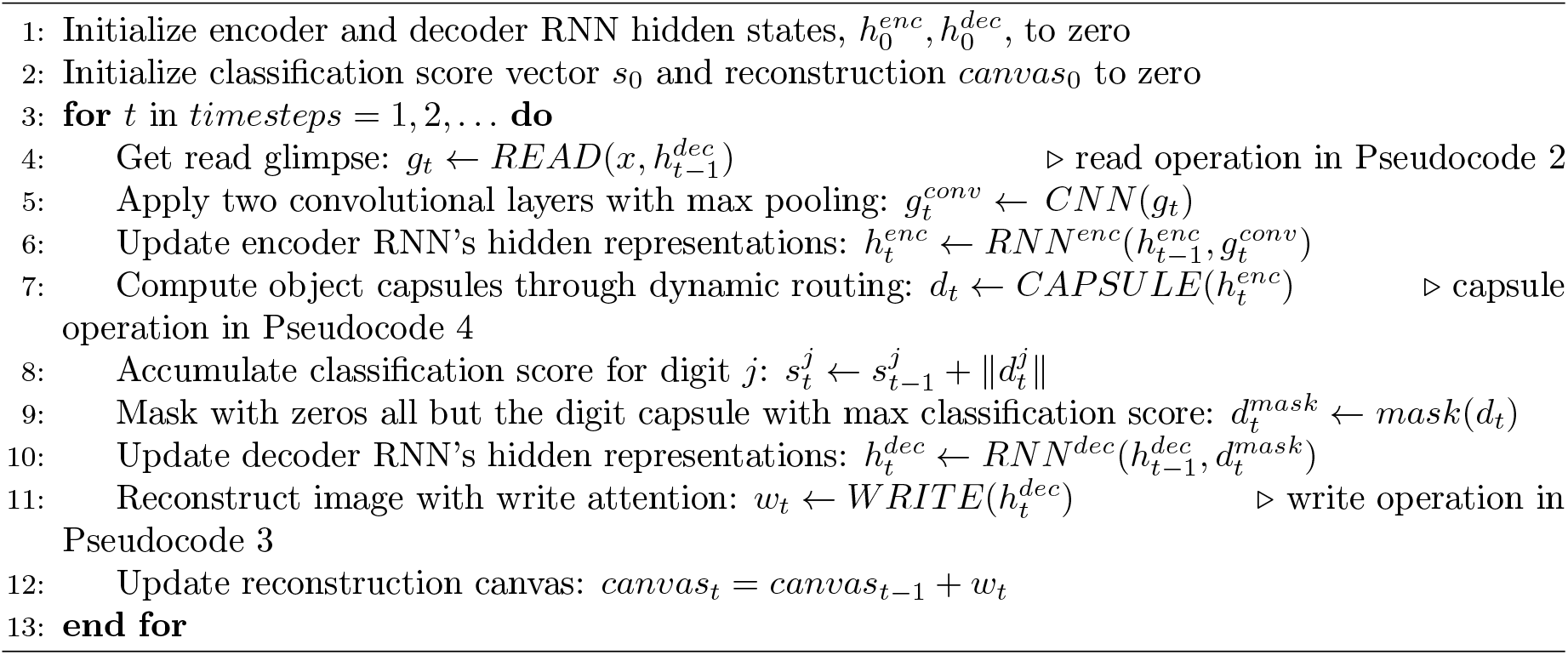

#### Read and Write Attention

At each timestep a glimpse, *g*_*t*_, is sampled through applying a grid of N *×* N Gaussian filters on the input image *x*. We set the glimpse size to 18 *×* 18 for our experiments, with a sample glimpse shown in Fig. 1 left. The Gaussian filters are generated using four parameters: *gX, gY, δ, σ*^2^, which specify the center coordinates of the attention window, the distance between equally spaced Gaussian filters in the grid, and the variance of the filters, respectively. All of these parameters are computed via a linear transformation of the previous step decoder RNN (Recurrent Neural Network) output 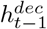 using a weight matrix *W*_*read*_, which makes the attention mechanism fully-differentiable. A similar procedure applies to the *write* attention operation. The decoder RNN output 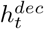 is linearly transformed into an M *×* M write patch *w*_*t*_ (set to 18 *×* 18 in our experiments), which is then multiplied by the Gaussian filters to reconstruct the written parts in the original image size (Fig. 1 right). The Gaussian filters used for the *write* operation differed from those used for the *read* operation, and were computed from four parameters obtained from a separate linear transformation, *W*_*write*_*attention*_, of the decoder RNN output 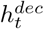. Detailed algorithms for Read and Write attention operations are provided in Pseudocode 2 and 3 in the Supplementary along with an illustration of the attention mechanisms (Fig. 5).

**Figure 5:**
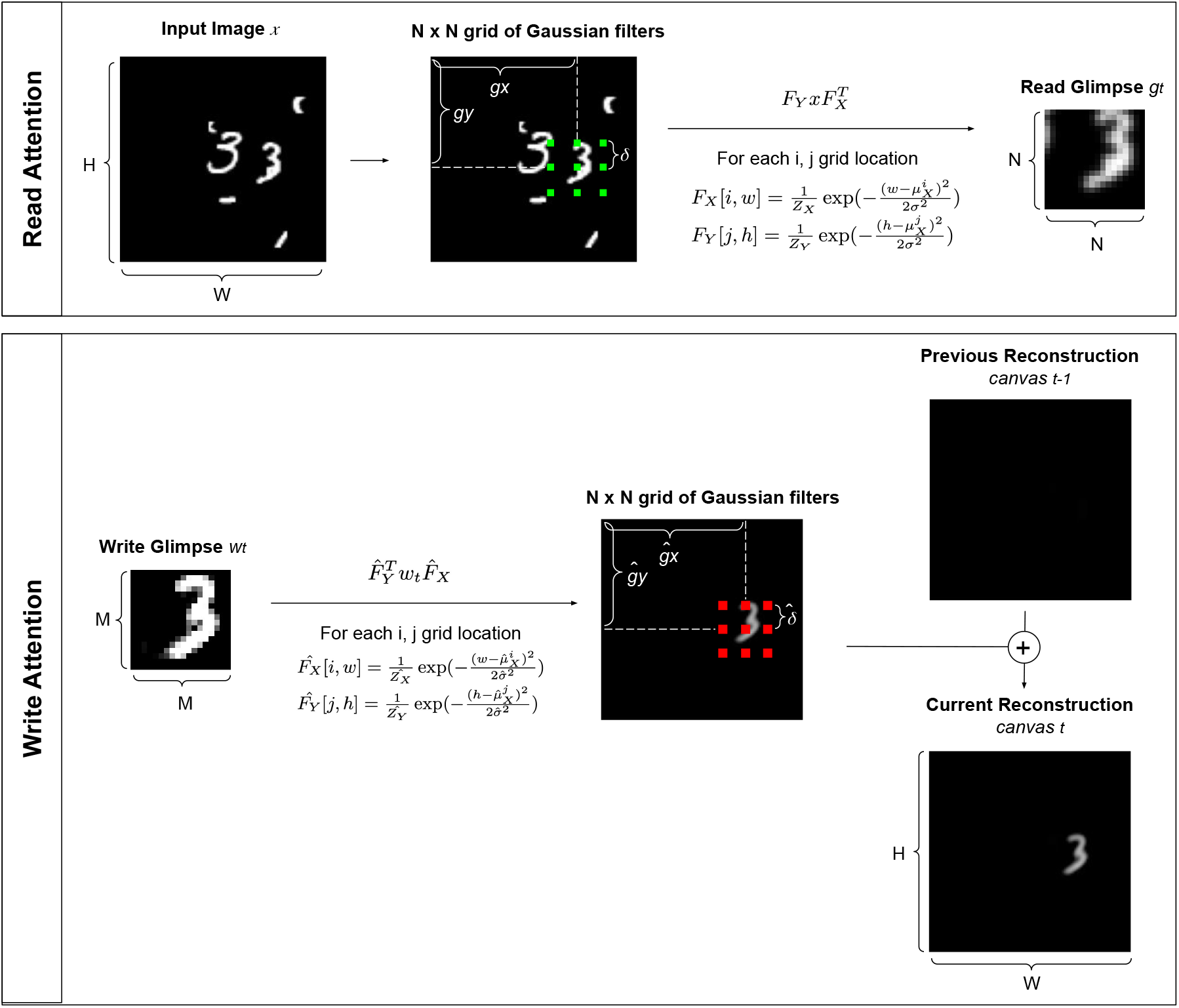
Visual illustrations of read and write attention mechanisms

##### Pseudocode 2

Read Attention

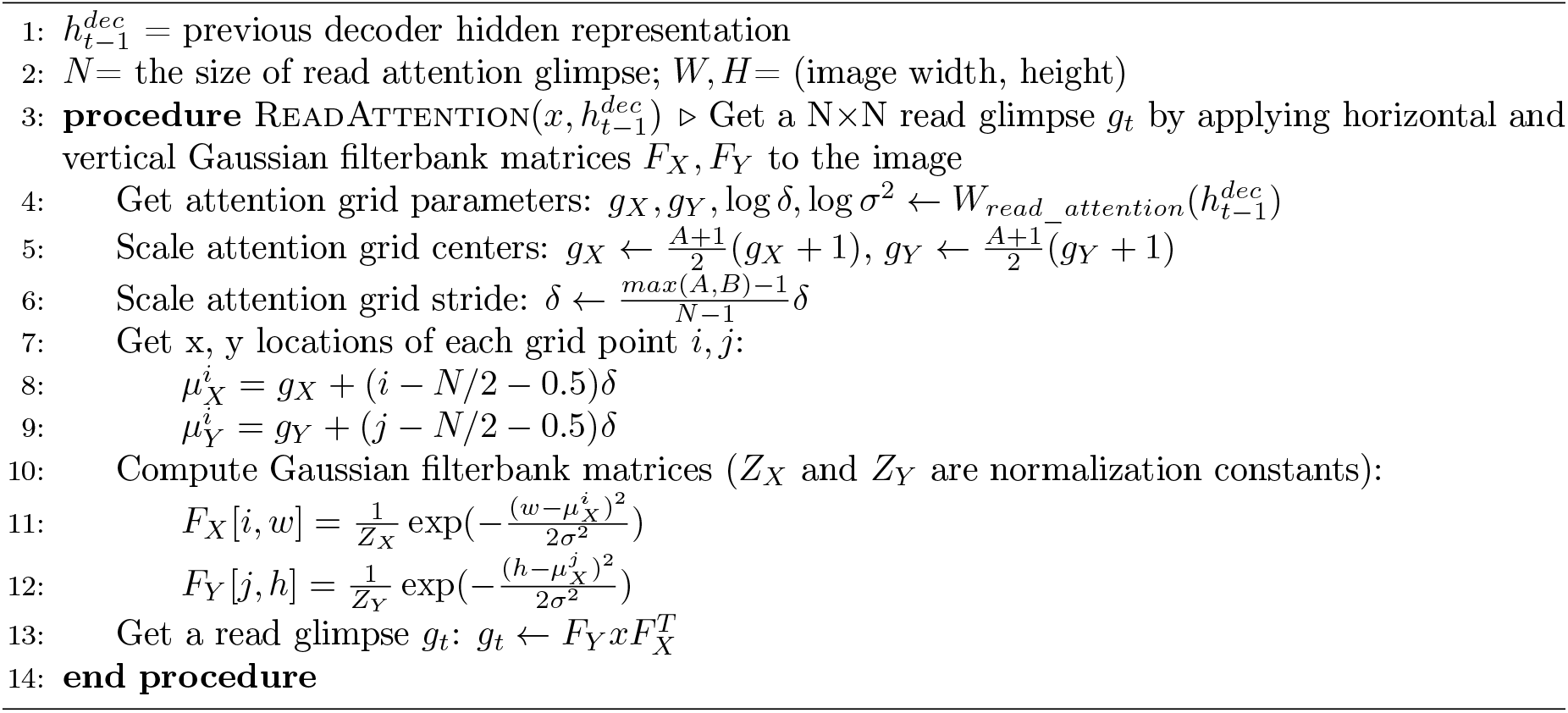

##### Pseudocode 3

Write Attention

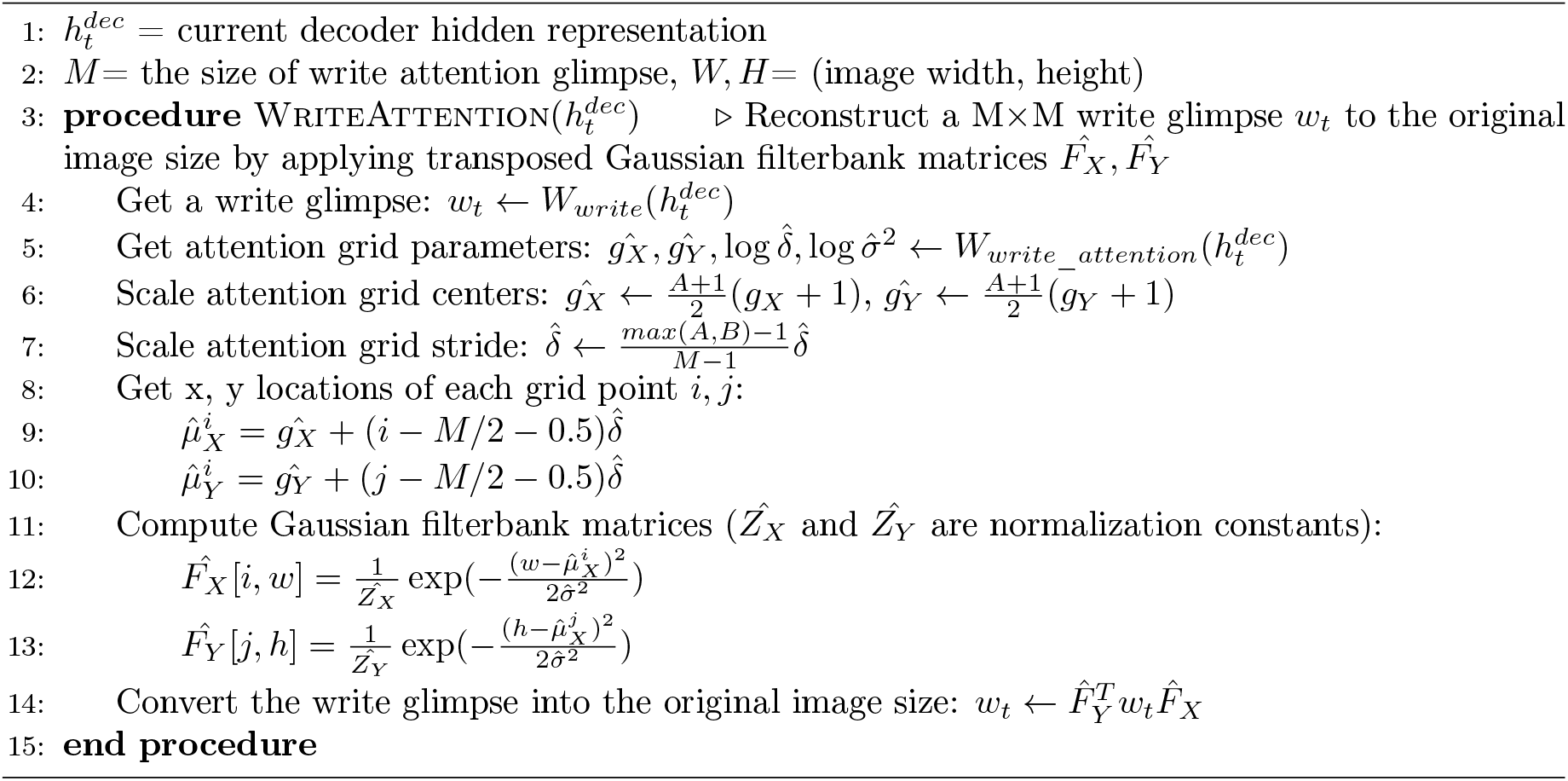

#### Encoder

After a glimpse is selected from the input image by the read attention operation, it is processed first using a two-layer convolutional neural network (CNN) with 32 filters in each layer. Kernel sizes were set to 5 and 3 respectively for the first and the second layers. Each convolutional layer is followed by max pooling with a kernel size of 2 and a stride of 2, and Rectified Linear Units (ReLU) were used for non-linear activation functions. Given the glimpse size of 18 *×* 18, the resulting 32 feature maps are of size 4 *×* 4. The feature maps, 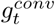, are reshaped (to a vector of size 512) and used as input to the encoder Recurrent Neural Network (RNN), along with the encoder RNN hidden state from the previous step; 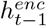. We used LSTM [36] units (size 512) for the recurrent layers in our model.

#### Latent Capsule Representations and Dynamic Routing

We use a vector implementation of capsules [66] where the length of the vector represents the existence of the visual entity and the orientation characterizes its visual properties. The primary level capsules are generated through a linear read out of the encoder RNN; 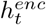. These capsules are meant to represent lower-level visual entities (“parts”) that belong to one of the higher-level capsules in the object capsule layer (“whole”). To find this part-whole relationship, we used the dynamic routing algorithm proposed by [66]. Dynamic routing is an iterative process where the assignments of parts to whole objects (coupling coefficients) are progressively determined by agreement between the two capsules (measured by the dot product between the two vector representations). Each primary level capsule (i) provides a prediction for each object level capsule (j). These predictions are then combined using the coupling coefficients (cij) to compute the object level capsule. Then the agreement (dot product) between the object level capsules and the predictions from each primary level capsule impacts the coupling coefficients for the next routing step.For example, if the prediction for a digit capsule *j* from a primary capsule *i*, 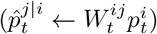, highly agrees with the computed digit capsule 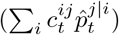, the coupling coefficient 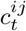 is enhanced so that more information is routed from primary capsule *i* to object capsule *j*. Coupling coefficients are normalized across the class capsule dimension following the max-min normalization [91] as in Eq. 1. Lower- and upper-bounds for normalization, *lb* and *ub*, were set to 0.01 and 1.0. This routing procedure iterates three times. We used this method instead of the softmax normalization in [66] because we observed the latter method would not differentiate between the coupling coefficients. In our experiments we used 40 primary level capsules, each a vector of size 8. The object capsules are vectors of size 16 and there are 10 of them corresponding to the 10 digit categories for the multi-object recognition task and 4 of them for the visual reasoning task. For the object level capsules, we use a squash function (the Supplementary Eq. 2) to ensure that its vector length is within the range of 0 to 1. For the mutli-object recognition task, these would represent the probability of a digit being present in the glimpse at each step. Once the routing is completed, we compute the vector length (L2 norm) of each object capsule to obtain classification scores. The final digit classification is predicted based on the scores accumulated over all timesteps. For the visual reasoning task, two capsules (among four total) were designated to be the response capsules and the cumulative length of these judgment capsules were used for same-different response. The Algorithms are provided in Pseudocode 4.

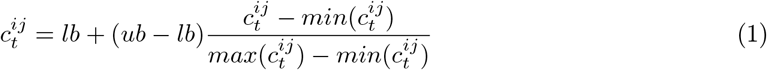

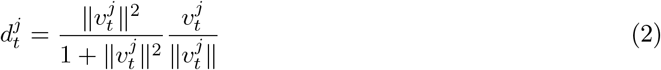

##### Pseudocode 4

Capsule Representation and Dynamic Routing

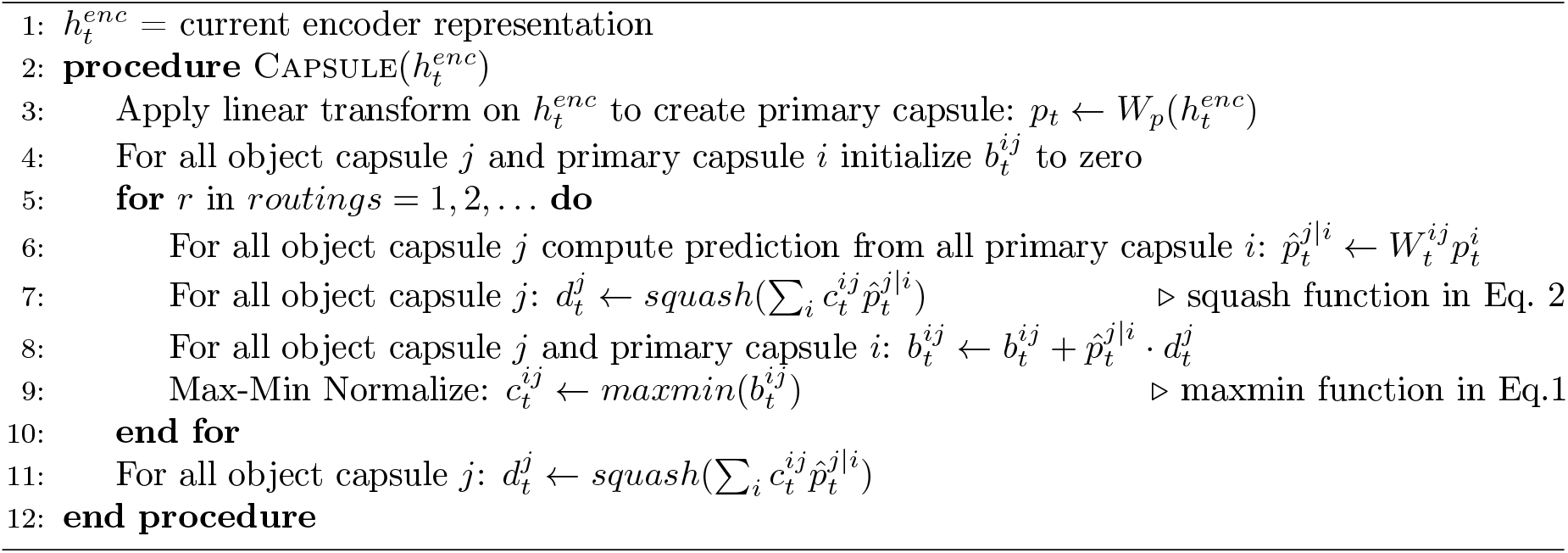

#### Decoder

The object capsules provide a structured representation that can be used for decoding and glimpse selection. We first mask the object capsules so that only the vector representation from the most active capsule is forwarded to the decoder RNN, which also inputs the hidden state from the previous step, 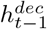. Because the decoder maintains through recurrence the ongoing and evolving object-based representation of the image, it is best suited to determine the next read glimpse location (as discussed earlier). The state of the decoder RNN is also used through two linear operations to determine what and where to write in the canvas to reconstruct the image.

### 4.3 Loss Function

OCRA outputs object classification scores (cumulative capsule lengths) and image reconstruction (cumulative write canvas). Losses are computed for each output and combined with a weighting as in Eq. 3. For reconstruction loss, we simply computed the mean squared differences in pixel intensities between the input image and the model’s reconstruction. For classification, we used margin loss (Eq. 4). For each class capsule j, the first term is only active if the target object is present (*T*_*j*_ *>* 0) where minimizing the loss pushes the capsule length to be bigger than target capsule length minus a small margin (m). The second term is only active when the target capsule length is zero and in that case minimizing the loss pushes the predicted capsule length to be below a small margin (m). For all the experiments in this paper we used Adam optimizer [44].

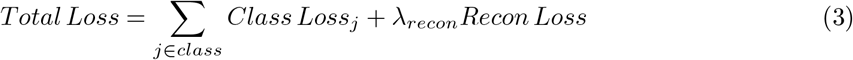

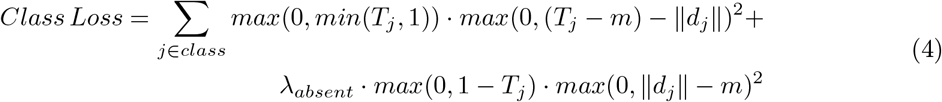

### 4.4 Measuring accuracy

When we convert the model scores to multi-object classification, we take into account the thresholds that are set in the loss function. If the model prediction for one class capsule is larger than 1.8 (conservatively selected to be slightly lower than 1.9), this signals the presence of two objects from this class in the image. If no class score is larger than this threshold, the top two highest scores are selected as the model predictions for the two objects in the image.

### 4.5 Training Regime and Hyperparameter selection

All the model training was done on a single GPU workstation. All the Models trained on the MultiMNIST task were stopped after 50 epochs of training (taking about 40 hours for the 10glimpse model). The models trained on the MultiMNIST-cluttered task were stopped after 1000 epochs (taking about 40 hours for the 5glimpse model). The models trained on the visual reasoning task task were stopped after 100 epochs (taking about 2 hours). We used the Adam optimizer [44] for all the experiments.

Table 4 provides all the hyperparameters used for the three tasks. The hyperparameters were mostly the same between the tasks with the few differences. For the MultiMNIST-cluttered task, we added one background capsule and utilized a reconstruction mask, both to allow the model to focus on the main objects and ignore the background clutter. The image is larger (100 *×* 100 vs 36 *×* 36) in the MultiMNIST-cluttered task, with most of it being empty, the reconstruction loss therefore has a much smaller range compared to the other task. For this reason, and the use of the reconstruction mask, we use a much larger weight for the reconstruction loss to make it comparable to the margin loss. For this task we added a background capsule to the 10 class capsules, similar in approach to [64]). This gives the model the choice to dynamically route background noise in its attention windows to a non-class capsule, thereby allowing the model to exclude the noise from its object representations.

**Table 4:**
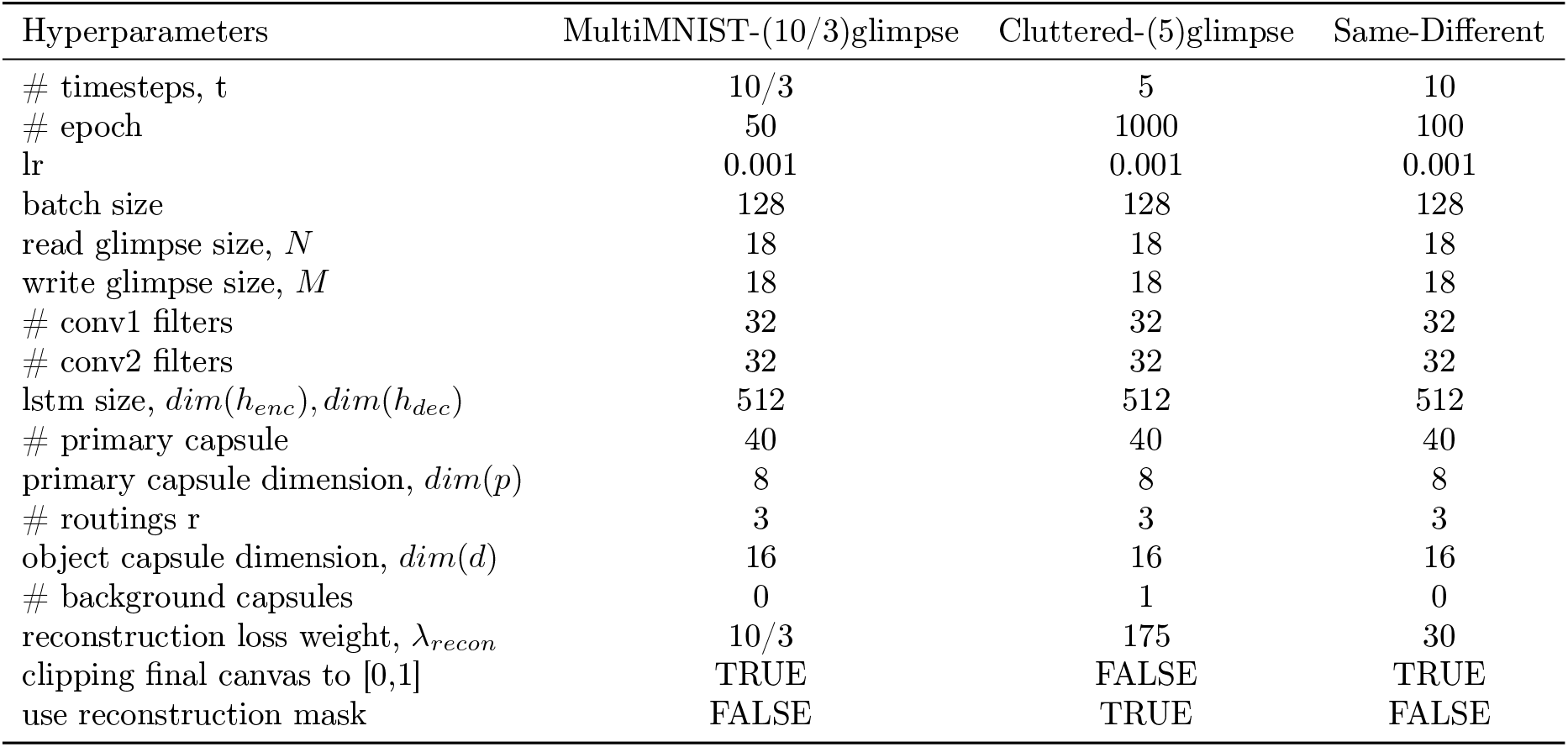
Hyperparameter Setting for MultiMNIST and MultiMNIST-cluttered tasks

| Hyperparameters | MultiMNIST-(10/3)glimpse | Cluttered-(5)glimpse | Same-Different |
| --- | --- | --- | --- |
| # timesteps, t | 10/3 | 5 | 10 |
| # epoch | 50 | 1000 | 100 |
| lr | 0.001 | 0.001 | 0.001 |
| batch size | 128 | 128 | 128 |
| read glimpse size, $N$ | 18 | 18 | 18 |
| write glimpse size, $M$ | 18 | 18 | 18 |
| # conv1 filters | 32 | 32 | 32 |
| # conv2 filters | 32 | 32 | 32 |
| lstm size, $\dim(h_{enc}), \dim(h_{dec})$ | 512 | 512 | 512 |
| # primary capsule | 40 | 40 | 40 |
| primary capsule dimension, $\dim(p)$ | 8 | 8 | 8 |
| # routings r | 3 | 3 | 3 |
| object capsule dimension, $\dim(d)$ | 16 | 16 | 16 |
| # background capsules | 0 | 1 | 0 |
| reconstruction loss weight, $\lambda_{recon}$ | 10/3 | 175 | 30 |
| clipping final canvas to [0,1] | TRUE | FALSE | TRUE |
| use reconstruction mask | FALSE | TRUE | FALSE |

For the MultiMNIST task, we clip the cumulative reconstruction canvas to be between [0,1] before comparing it to the input image. This allows the model to overlap different segments in the multi-step process of writing to the canvas without increasing the loss, improving model reconstruction given the high degree of overlap between the digits.

For the Visual Reasoning task, we used mostly similar hyperparameters. The number of class capsules were set to 4 as explained in the main text.

## A Supplementary Material

**Figure S1:**
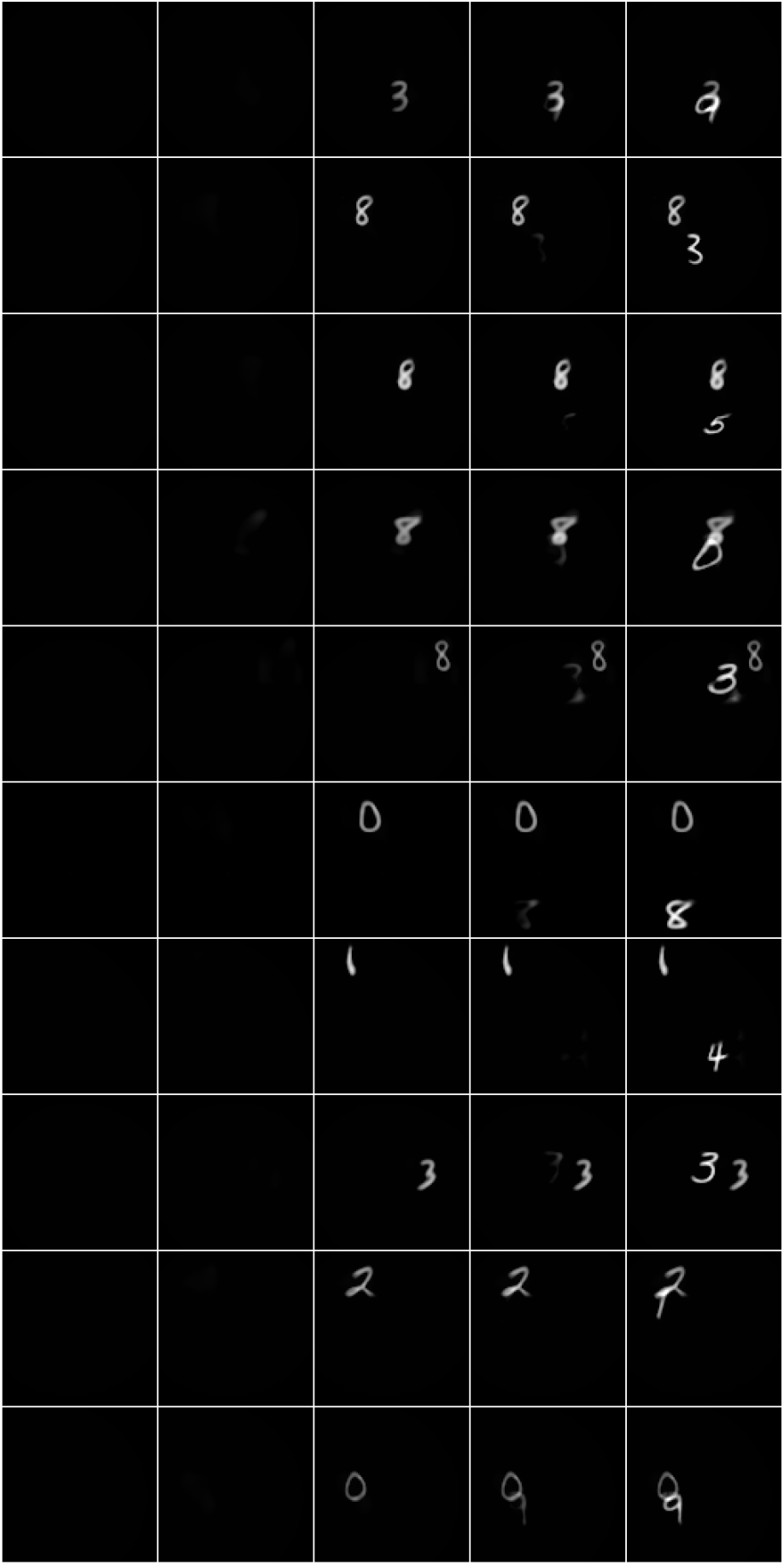
OCRA MultiMNIST-cluttered output with 5 glimpses showing the gradual object-based reconstruction on the cumulative canvas.

**Figure S2:**
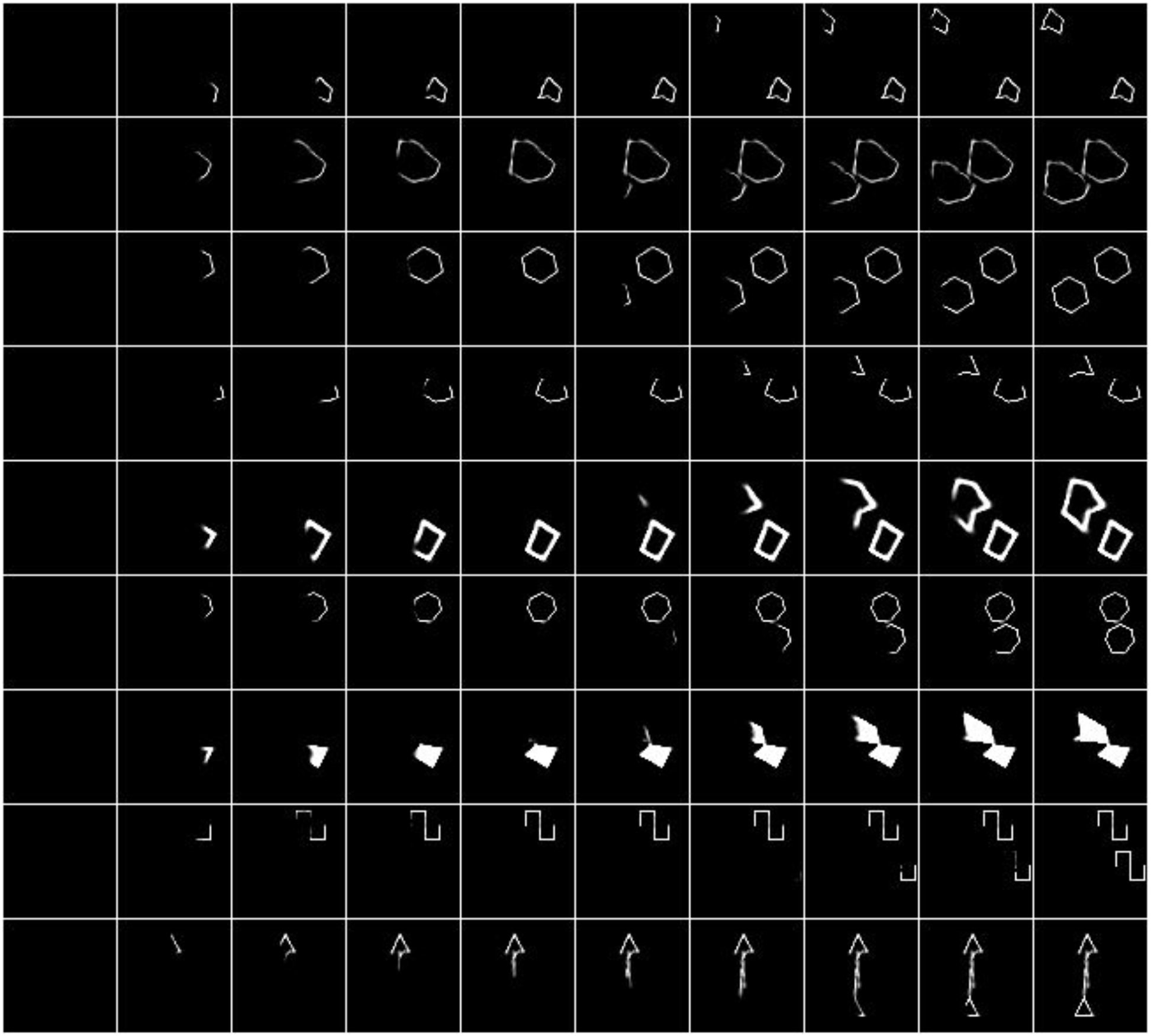
OCRA visual reasoning with 10 glimpses showing the gradual object-based reconstruction on the cumulative canvas.

